# Towards understanding cancer stem cell heterogeneity in the tumor microenvironment

**DOI:** 10.1101/408823

**Authors:** Federico Bocci, Larisa Gearhart-Serna, Marcelo Boareto, Mariana Ribeiro, Eshel Ben-Jacob, Gayathri R. Devi, Herbert Levine, José Nelson Onuchic, Mohit Kumar Jolly

## Abstract

The Epithelial-Mesenchymal Transition (EMT) and Cancer Stem Cell (CSC) formation are two paramount processes driving tumor progression, therapy resistance and cancer metastasis. Some recent experiments show that cells with varying EMT and CSC phenotypes are spatially segregated in the primary tumor. The underlying mechanisms generating such spatiotemporal dynamics and heterogeneity in the tumor micro-environment, however, remain largely unexplored. Here, we show through a mechanism-based dynamical model that the diffusion of EMT-inducing signals such as TGF-β in a tumor tissue, together with non-cell autonomous control of EMT and CSC decision-making via juxtacrine signaling mediated via the Notch signaling pathway, can explain experimentally observed disparate localization of subsets of CSCs with varying EMT states in the tumor. Our simulations show that the more mesenchymal CSCs lie at the invasive edge, while the hybrid epithelial/mesenchymal (E/M) CSCs reside in the tumor interior. Further, motivated by the role of Notch-Jagged signaling in mediating EMT and stemness, we investigated the microenvironmental factors that promote Notch-Jagged signaling. We show that many inflammatory cytokines that can promote Notch-Jagged signaling such as IL-6 can (a) stabilize a hybrid E/M phenotype, (b) increase the likelihood of spatial proximity of hybrid E/M cells, and (c) expand the fraction of CSCs. To validate the predicted connection between Notch-Jagged signaling and stemness, we knocked down JAG1 in hybrid E/M SUM149 human breast cancer cells *in vitro*. JAG1 knockdown significantly restricted organoid formation, confirming the key role that Notch-Jagged signaling can play in tumor progression. Together, our integrated computational-experimental framework reveals the underlying principles of spatiotemporal dynamics of EMT and CSCs in the tumor microenvironment.

**Significance statement:** The presence of heterogeneous subsets of cancer stem cells (CSCs) remains a clinical challenge. These subsets often occupy different regions in the primary tumor and have varied epithelial-mesenchymal phenotypes. Here, we device a theoretical framework to investigate how the tumor microenvironment (TME) modulates this spatial patterning. We find that a spatial gradient of EMT-inducing signal, coupled with juxtacrine Notch-JAG1 signaling triggered by inflammatory cytokines in TME, explains this spatial heterogeneity. Finally, *in vitro* JAG1 knockdown experiments in triple negative breast cancer cells severely restricts the growth of tumor organoid, hence validating the association between JAG1 and CSC fraction. Our results offer insights into principles of spatiotemporal patterning in TME, and identifies a relevant target to alleviate multiple CSC subsets – JAG1.

## Introduction

The tumor microenvironment (TME) offers intriguing questions about the spatiotemporal dynamics of pattern formation. Abundant phenotypic and functional heterogeneity (1–3), coupled with varying concentrations of nutrient availability (4) and bidirectional crosstalk among constituent cells (5), can lead to the formation of complex patterns correlated with aggressive pathological behaviors. Two interconnected hallmarks of cellular plasticity that contribute to this spatiotemporal heterogeneity are the Epithelial-Mesenchymal Transition (EMT) and the generation of Cancer Stem Cells (CSCs). These are a ‘dangerous duo’ that can cooperatively promote tumor progression, metastasis, therapy resistance and tumor relapse (6). Remarkably, the initial proposition of the involvement of EMT in metastasis was based on spatially varying levels of β-catenin (7), where cells at the invasive edge of the tumor and those in central areas contained varying subcellular localization of β-catenin and E-cadherin, thus affecting cell adhesion and migration. Extensive characterization since then has identified that the activation of EMT and the localization of different subsets of CSCs is spatially quite heterogeneous within a tumor (8, 9). This heterogeneity can arise from bidirectional interplay among tumor cells and stromal cells, thus highlighting the contributions that the non-cell autonomous effects in the activation of EMT and CSCs can have on spatiotemporal dynamics in the TME.

These non-cell autonomous effects can be modulated by cell-cell communication pathways that crosstalk with intracellular signaling networks governing EMT and CSCs. One canonical case is the Notch-Jagged signaling pathway, which has been implicated in metastasis, drug resistance and tumor relapse (10). Notch-Jagged signaling has been studied in multiple developmental contexts for its role in cellular decision-making and tissue patterning (11–13). Recent *in silico*, *in vitro* and *in vivo* observations have suggested that Notch-Jagged signaling can promote a hybrid epithelial /mesenchymal (E/M) phenotype and traits similar to CSCs (5, 14–16). Moreover, Notch-Jagged signaling among cancer cells can facilitate the formation of clusters of circulating tumor cells (CTCs) – the ‘bad actors’ of cancer metastasis (15–17). Furthermore, Notch-Jagged signaling between cancer cells and stromal cells and/or cells in metastatic niche can aggravate tumor progression (18). Put together, hybrid E/M cells with CSC-like traits and activated Notch-Jagged signaling perhaps represents the ‘fittest’ phenotype for metastasis (19–22).

Several factors in the tumor microenvironment can activate the Notch-Jagged pathway. For instance, many inflammatory cytokines such as IL-6, IL-1β and TNF-α can promote Notch-Jagged signaling by increasing the intracellular production of Jagged. Some of them can also decrease the production of Delta, a competing ligand that binds to the Notch receptor (23–26). Inflammation is a hallmark of damaged tissues: acute inflammation plays a crucial role in wound healing; chronic inflammation predisposes the damaged tissue to the development of cancer. Also, cancer patients treated with chemotherapy and radiation have elevated inflammation (27, 28), often via the abovementioned inflammatory cytokines that can aggravate tumor progression by both promoting EMT and CSC formation (29, 30). The operating principles behind the complex spatiotemporal interplay between signaling cues diffusing in the TME, non-cell autonomous signaling via Notch-Jagged, and the intra-cellular molecular machinery regulating EMT and CSC, however, remain largely elusive.

Here, we propose a mathematical modeling framework that captures the experimentally identified interconnections among inflammatory cytokines, EMT, CSCs, and Notch signaling and decodes the emergent spatial patterns of different subsets of CSCs with different EMT phenotypes. First, we show that the production and diffusion of an EMT-inducing signal (such as TGF-β), together with Notch signaling, can robustly recapitulate experimental observations that while mesenchymal CSCs are localized towards the tumor-stroma periphery, hybrid E/M CSCs are typically located in the tumor interior (9). Further, modeling the effect of inflammatory cytokines in the TME shows that these cytokines can stabilize a hybrid E/M phenotype and increases the frequency of CSCs by activating Notch-Jagged signaling. We experimentally validate the functional relation between Notch-Jagged signaling and stemness by knocking down JAG1 in hybrid E/M SUM149 inflammatory breast cancer cells and showing a significant impairment of tumor formation *in vitro*. Put together, our model yields valuable insights into the interconnected spatiotemporal dynamics of EMT, CSCs, and Notch-Jagged signaling, identifies the mechanisms underlying experimental observations, and generates testable predictions that have been validated in SUM149 breast cancer cells.

## Results

### A gradient of EMT-inducing signal recapitulates the experimentally observed spatial segregation of different EMT phenotypes

The interplay of different micro-environmental signals can give rise to intricate spatial arrangements of cells with different EMT phenotypes and/or different subsets of CSCs. Immunofluorescence analysis of breast carcinoma tissue revealed that different populations of breast CSCs with different EMT profiles can localize in anatomically distinct regions of a tumor (Fig. 1A) (9). A CD24^-^CD44^+^ mesenchymal-like breast cancer stem cell (BCSC) population was localized at the invasive edge of tumor, while an ALDH1^+^ epithelial-like BCSC population was localized in the interior close to the tumor stroma (Fig. 1A). Follow-up studies have characterized these ALDH1+ cells as showing a hybrid E/M, and not a purely epithelial, signature (31).

**Figure 1.**
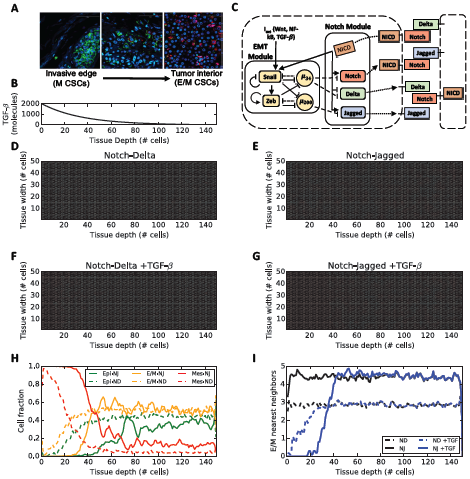
EMT phenotype patterning in the presence of EMT-induced Notch signaling. **(A)** Immunofluorescent staining of CD24 (magenta), CD44 (green), ALDH1 (red) and DAPI (blue) in human invasive breast carcinoma adapted from Liu, Cong et al. (9) (in agreement with the journal copyright guidelines). **(B)** Coupling between Notch pathway and core EMT regulatory circuit. Solid arrows/bars represent transcriptional activation/inhibition, while dashed lines represent post-translational inhibition by micro-RNAs. Dotted lines represent translocation of Notch, Delta, and Jagged to the cell surface. **(C)** Concentration of EMT-inducing signal as a function of tissue depth in the layer. Because the signal is secreted uniformly at one end of the layer, the profile is constant across the layer width. **(D)** EMT phenotype distribution in the cell layer after 120 hours of equilibration starting from randomized initial conditions and for a strong Notch-Delta signaling (g_D_ = 90 molec/h, g_J_ = 20 molec/h). Green, yellow and red colors denote epithelial (E), hybrid (E/M) and mesenchymal (M) cells, respectively. **(E)** Same as panel (D) for strong Notch-Jagged signaling (g_D_ = 20 molec/h, g_J_ = 50 molec/h) **(F)** Same as panel (D) in the presence of TGF-β gradient in the tissue layer. **(G)** Same as panel (E) in the presence of TGF-β gradient in the tissue layer. **(H)** Fraction of E, hybrid E/M and M cells as a function of tissue depth corresponding to panel (F) (dashed lines) and (G) (continuous lines). **(I)** Average number of E/M nearest neighbors of the hybrid E/M cells as a function of tissue depth for the 4 cases of panels (D)-(E)-(F)-(G). (H) and (I) present an average over 10 simulations starting from random initial conditions.

To decipher the signaling mechanisms that may underlie such heterogeneous distribution of EMT phenotypes, we extended our previously developed mathematical model that couples a core EMT regulatory circuit with the juxtacrine Notch signaling pathway (5). Here, we consider the effect of a diffusing EMT-inducing signaling (such as TGF-β) on our multi-cell lattice setup consisting of (50*150 cells). A spatial gradient of TGF-β diffuses from one end of the layer (the invasive edge of tumor, as shown in Fig. 1A) towards the tumor interior, mimicking the secretion of TGF-β by stromal cells (32) (Fig 1B). The width/depth ration of the lattice was chosen to evaluate the effect of gradients of signaling molecules in different tissue areas that represent anatomically distinct regions of a tumor.

The Notch signaling pathway is activated when one of its transmembrane ligands (Delta, Jagged) binds to the transmembrane receptor Notch belonging to a neighboring cell, leading to the release of NICD (Notch Intracellular Domain). NICD regulates several target genes resulting in the activation of Notch and Jagged and the inhibition of Delta (33) (Fig. 1C, Notch module). Therefore, Notch-Jagged signaling between two cells enables a similar cell fate (‘lateral induction’) where both cells can send and receive signals, due to the presence of ligand (Jagged) and receptor (Notch), thus called a hybrid Sender/Receiver phenotype. On the other hand, Notch-Delta signaling promotes opposite cell fates (‘lateral inhibition’) where one cell has high Notch (thus, acting as a Receptor) and the other cell has high Delta (thus, behaving as a Sender) (34, 35). In our model, Jagged refers to JAG1 that has been found to be particularly involved in lateral induction as well as tumor progression (15, 18, 36). Further, NICD activates EMT by activating the EMT-inducing transcription factor (EMT-TF) SNAIL (5) (Fig. 1C). A core regulatory circuit for EMT is comprised of two families of EMT-TFs (SNAIL, ZEB) and two families of EMT-inhibiting microRNAs (miR-34, miR-200) that mutually inhibit each other (37). Additional inputs from pathways such as TGF-β can activate EMT (37) (Fig. 1C, EMT module). This EMT circuit can enable three cell phenotypes: epithelial (high microRNAs, low EMT-TFs), hybrid E/M (intermediate microRNAs, intermediate EMT-TFs), and mesenchymal (low microRNAs, high EMT-TFs) (37). These microRNAs can inhibit the activation of Notch signaling (5); thus, the activation of EMT can also lead to activated Notch signaling.

In the absence of a TGF-β gradient, Notch-Jagged signaling generates spatial patterns with clusters of hybrid E/M cells, but Notch-Delta signaling leads to an alternating arrangement of epithelial and hybrid E/M cells, similar to our previous results (5, 38) (Fig. 1D-E and movies M1, M2). Introducing the gradient of the EMT-inducing signal TGF-β through the tissue, however, generates spatial segregation of different EMT phenotypes. Cells close to the invasive edge, where TGF-β is secreted, undergo a complete EMT while cells in the interior, at low TGF-β exposure, are mostly epithelial and hybrid E/M (Fig. 1F-G and movies M3, M4). Specifically, the fraction of hybrid E/M cells is similar in the Notch-Delta and Notch-Jagged cases (Fig. 1H and S1), but the patterning is remarkably different. Notch-Jagged signaling, but not Notch-Delta signaling, enables a pattern where the hybrid E/M cells are much more likely to be surrounded by other hybrid E/M cells; thus, leading to the formation of clusters of hybrid E/M cells, in the tissue interior (Fig. 1I). JAG1 has been observed to be among top differentially expressed genes in clusters of CTCs (16), thus supporting the prediction of our model.

Therefore, the secretion and diffusion of EMT-inducing signals such as TGF-β from the tumor-stroma boundary can generate the spatial segregation of cells with different EMT phenotypes experimentally observed in carcinomas (7, 9) in either scenarios – when cells are communicating via Notch-Delta or Notch-Jagged. Furthermore, experimental results indicate that the generation of CSCs – a process intricately tied with EMT (39, 40) - can be produced both at the tumor edge (mesenchymal CSCs) and in the interior region at lower exposure to EMT-inducer (hybrid E/M CSCs). Thus, a more complete characterization of the tumor tissue requires understanding the signaling mechanisms in the TME that can mediate CSC properties.

### Inflammatory cytokines can promote a hybrid E/M phenotype by increasing Notch-Jagged signaling

Previous observations have suggested a key role of the Notch-Jagged signaling axis in mediating CSC properties (14, 41–43). Therefore, we next focused on what signals in the TME can promote Notch-Jagged signaling. Inflammatory cytokines such as IL-6, IL-1β, TNF-α and the pathways they involve (such as NF-?B) can enhance the production of Jagged and, in some cases, also inhibit that of Delta (23–26). To investigate the effect of these cytokines, we introduced an external variable (*C_EXT_*) that acts as an activator of Jagged and an inhibitor of Delta (see Methods). We vary *C_EXT_* to characterize the effect of different inhibition/activation strengths arising in a concentration-specific or cytokine-specific way.

As a first step towards understanding the effect of inflammatory cytokines on Notch signaling and the plasticity of tumor cells, we analyzed the dynamics of an individual cell that is exposed to variable levels of inflammatory cytokines (*C_EXT_*) and Jagged ligands (*J_EXT_*) that can bind to the Notch receptor present at the cell surface and induce the EMT regulatory cascade.

We first plotted the levels of miR-200 (gatekeepers of epithelial phenotype (37, 44)) upon full equilibration of the system, as a function of the external ligand available *(J_EXT_*), under conditions of low levels of inflammatory cytokines (*C_EXT_* = 1000 molecules). The cell is initially in an epithelial (E) phenotype (high levels of miR-200), and exhibits a ‘Sender’ (S) Notch state characterized by a low expression of Notch receptor and a high expression of ligand Delta (Fig 2A - (E), (R)). For increasing values of *J_EXT_*, Notch signaling is activated, triggering a shift to a ‘Receiver’ Notch (R) state characterized by a high expression of Notch receptor and a low expression of ligand Delta. EMT, however, is not yet triggered (green shaded region in Fig 2A - (E), (R)). A larger level of *J_EXT_* further activates Notch signaling and induces a partial EMT, or a transition to a hybrid E/M phenotype. Concomitantly, intracellular Jagged production is also elevated as the inhibition of Jagged by miR-200 is relieved. Thus, the cell attains a hybrid Sender/Receiver (S/R) Notch state – (orange shaded region in Fig 2A – (E/M), (S/R)). A further increase in *J_EXT_* induces a stronger activation of the EMT circuitry, driving the cells towards a mesenchymal state (red shaded region in Fig 2A – (M), (S/R)). It is worth pointing out that our model operates in a parameter regime where the Notch signaling is functionally active in epithelial cells. It is possible, however, that the epithelial cells have ‘inactive’ Notch signaling (Fig. S2), as seen in some contexts.

**Figure 2.**
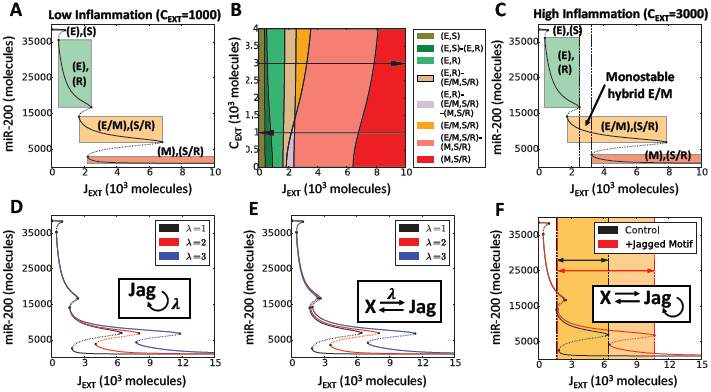
Inflammation stabilizes a hybrid E/M phenotype. **(A)** Bifurcation curves of miR-200 as a function of J_EXT_ for low inflammation (C_ext_ = 1000 molecules). **(B)** Cell phenotype diagram as a function of external Jagged (J_EXT_) and external cytokines (C_EXT_). The different colors represent portions of parameter space characterized by monostability or multistability of the coupled Notch-EMT system. **(C)** represents the same case as (A), but for high inflammation (C_ext_ = 3000 molecules). Solid lines represent stable steady states, and dotted lines represent unstable steady states. Vertical dotted lines in (C) depict the range of control parameter values that allows for monostability of the (E/M, S/R) state. The colored rectangles in (A) and (C) elucidate the interval of (J_EXT_) values for which different states of coupled EMT-Notch circuitry are stable, and the corresponding level of miR-200. For this simulation, the external concentration of Notch and Delta are fixed at N_EXT_ = 10000 molecules, D_EXT_ = 0 molecules (35). Bifurcation diagrams for all model’s variables are presented in Fig. S3B. **(D-F)** Bifurcation diagram of miR-200 in presence of self-activation of Jagged (D), positive feedback loop between Jagged and another additional component (E) and a combination of both (F). Hill coefficient(s) is(are), unless stated otherwise, n = 2. In (D), λ is the fold change in production rate of Jagged due to the activation by X, while in (E) it represents the fold-change of both interactions. In (F), all λ = 2.

Next, to better understand the role of inflammatory cytokines in mediating this bifurcation diagram, we plotted a two-dimensional phenotype diagram, varying the levels of both *C_EXT_* and *J_EXT_* (Fig. 2B). This diagram displays multiple phases (i.e. sets of co-existing phenotypes under the same set of physiological conditions). The population distribution, when cells can attain more than one phenotype, depends on the local microenvironment, and the varied genetic and epigenetic landscapes of individual cells. Remarkably, at low levels of inflammatory cytokines, the hybrid (S/R, E/M) state is found only in multi-stable regions, i.e. only in combination with other possible cell states (Fig. 2B). Conversely, this cell state can exist by itself (monostable phase) when the cytokine levels are high. This remarkable difference can be visualized through a bifurcation diagram of miR-200 for a case of high levels of inflammatory cytokines (*C_EXT_* = 3000 molecules), where the region of stability of a hybrid E/M phenotype significantly increases (shown by dotted rectangle in Fig. 2C). A similar effect is observed in a single cell model driven by the external ligand Delta (Fig. S3A-C).

The effect of inflammatory cytokines on Notch signaling can often be mediated by the inflammatory response transcription factor NF-kB. For instance, NF-kB governs the effect of TNF-α on Jagged levels (23). Further, Jagged can activate NF-kB (45, 46), thus indirectly leading to its self-activation. Also, IL-6/IL-6R signaling can be promoted via autocrine effects mediated through EMT-Notch circuitry (47). To assess the effect of such amplification and/or autocrine signaling, we tested various circuit motifs that result in Jagged overexpression, by computing the bifurcation diagram of miR-200. Our motifs include Jagged self-activation (Fig. 2D), mutual positive activation (Fig. 2E) and the combination of these two possibilities (Fig. 2F). All motifs contributed to further stabilizing a hybrid E/M phenotype (Fig. 2D-F). These results suggest that autocrine and/or paracrine effects of inflammatory cytokines capable of upregulating Notch-Jagged signaling, such as IL-6, can stabilize a hybrid E/M phenotype.

Generalizing the model to a multi-cell scenario, these cytokines increase both the fraction of E/M cells in the tissue (Figure S4A-B) and the likelihood of physical contact between these hybrid E/M cells (Fig. S4C) when the signaling is dominated by Notch-Jagged signaling, but not by Notch-Delta signaling (Fig. S5), reminiscent of what is observed in Fig. 1G-I.

Together, these results indicate that inflammation can expand the range of physiological conditions for which cells can adopt a hybrid state in terms of cellular plasticity (E/M) and inter-cellular signaling (S/R) by activating Notch-Jagged signaling. This behavior is reminiscent of previous observations that JAG1 can act as a ‘phenotypic stability factor’ for a hybrid E/M phenotype (5).

## Inflammatory cytokines increase the CSC population by enhancing Notch-Jagged signaling

We have discussed how the gradients of EMT-inducing signals, together with Notch-Jagged signaling promoted by inflammatory cytokines such as IL-6, can enable a subpopulation of cancer cells to exhibit a hybrid E/M phenotype. A hybrid E/M phenotype has been proposed to often possess the typical properties of cancer stem cells (CSCs) (19–22, 48). Furthermore, recent experiments showed that various inflammatory cytokines increase the CSC population in a tumor (49). Thus, we explore the effect of cytokines on stemness. Within our framework, we define stemness based on activation of Notch-Jagged signaling (Figure S6A). Therefore, a cell is classified as CSC if it has high levels of NICD and Jagged. This classification is based on observations in multiple cancer types such as glioblastoma, pancreatic cancer, colon cancer and basal-like breast cancer that Notch and Jagged are overexpressed in CSCs as compared to non-CSCs (50–53). Further, our previous *in silico* results indicated that enhanced Notch-Jagged signaling is likely to correlate with the acquisition of stem-like traits (14). Thus, to elucidate the connection between inflammation, EMT and CSCs, we determined the frequency of CSCs in a population of a (100*100) multi-cell lattice. We chose a square lattice because no spatial gradient was applied in this simulation.

To probe the variation in CSC population due to inflammatory cytokines, we set up a simulation where the multi-cell layer is exposed to these cytokines for a variable amount of time (t_i_) and then allowed to equilibrate (Figure 3A). The fraction of CSC initially increases sharply as soon as inflammation is applied. Afterwards, if the inflammation is maintained for long enough, the conversion rate of non-CSC to CSC diminishes. Once the inflammatory stimulus is removed, the CSC fraction decreases (Figure 3B). Interestingly, this rate of return to initial state depends on the length of the inflammation period. If inflammation is applied for a short period, the fraction of CSCs in the tissue quickly returns to the initial value (i.e. corresponding fraction of CSC with no applied inflammation). Conversely, compared to the control value, chronic inflammation maintains a significantly higher CSC population for a longer period after the inflammation is removed (compare the “4 h” and “16 h” curves with the “64 h” and “128 h” curves in Figure 3B, and Fig. S6B). This observation suggests that the effects of inflammation on a tissue can be a long-lasting phenomenon and prevail for a significant time after the stimulus is removed.

**Figure 3.**
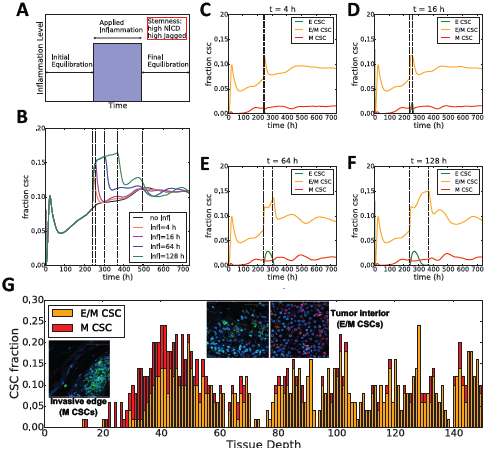
Inflammation increases the CSC population. **A)** Schematic of simulation setup: the two-dimensional layer of cells undergoes initial equilibration for 240 hours (initial conditions for proteins and micro-RNAs are extracted randomly); an inflammatory signal constant through the layer (C_ext_ = 3000 molecules) is applied for a variable time interval (blue region); after the inflammation is removed, the system equilibrates. **B)** Temporal dynamics of the fraction of CSC for different durations of applied inflammation and comparison to control (no applied inflammation, black curve). The first vertical dotted line from left indicates the time when the inflammation is applied (same for all curves); the four successive dotted lines depict the end of the applied inflammation for the different curves. **C-F)** Temporal dynamics of the fraction of epithelial, hybrid E/M and mesenchymal CSC. In C), for a short inflammation period (t=4 hours) the spike in CSC population is due to hybrid E/M cells. In this simulation, the production rates of Jagged and Delta are g_J_ = 50 molecules/h, g_D_ = 25 molecules/h, respectively (as in Fig. 3). **(G)** Fraction of CSCs with different EMT phenotypes as a function of tissue depth in the cell layer for the (Notch-Jagged, TGF-β gradient) of panel Fig. 1G. Insets show CSC the spatial distribution of M-CSC by the invasive edge of the tumor and E/M-CSC in the tumor interior as reported by Liu, Cong et al. (9).

Specifically, the initial spike in CSC fraction in response to inflammation is essentially a conversion of hybrid E/M non-CSCs into hybrid E/M CSCs (Figure 3C-F, orange curves, and Fig. S6C). In other words, these are hybrid E/M cells that had relatively low levels of either NICD or Jagged and hence were not classified as CSCs before the inflammatory stimulus was applied. After the initial spike, the fraction of hybrid E/M CSC remains nearly constant, while the fraction of epithelial CSCs increases (Figure 3D, E, green curves, and Fig. S6C). If the inflammation is maintained for longer time, epithelial CSCs undergo partial EMT and become hybrid E/M CSCs (see the reduction of epithelial CSCs and the corresponding increase in hybrid E/M CSCs in Figure 3F). Interestingly, inflammation does not alter the fraction of mesenchymal CSCs (Figure 3C-F, red curves, and Fig. S6C). Overall, these results suggest a strong correlation between the cytokine-induced hybrid E/M phenotype and the acquisition of stem-like traits.

Finally, the quantification of stemness based on Notch-Jagged signaling can explain the segregation of CSCs with different EMT phenotypes as observed experimentally (9) in the case of a tissue layer exposed to TGF-β gradient and strong Notch-Jagged signaling. Specifically, our model predicts a front of mesenchymal CSC toward the tumor-stroma interface, where the exposure to TGF-β is high, and a high fraction of hybrid E/M CSC in tumor interior (Fig. 3G and bottom panel of movies M1-4), in excellent agreement with experimental data (panels in Fig. 3G). This prediction is in good agreement with our previous observation that an E/M-CSC phenotype can switch to a M-CSC phenotype upon activation of TGF-β downstream targets (14). These results offer a unifying framework for the concepts of intratumoral heterogeneity, EMT and CSCs, and indicate the crucial role of JAG1 in simultaneously promoting a hybrid E/M phenotype and the acquisition of stem properties.

### JAG1 knockdown restricts emboli formation in inflammatory breast cancer cells SUM149

To validate our prediction that enhanced Notch-Jagged signaling can enhance the acquisition of a stem-like, more aggressive phenotype, we investigated the proliferation and tumor formation propensity in a model of triple negative, inflammatory breast cancer (IBC) cell line SUM149 in response to knockdown of JAG1. SUM149 cells constitutively express epidermal growth factor receptor and co-express E-cadherin and N-cadherin, and have been previously characterized as exhibiting a hybrid E/M phenotype (54–56). Further, flow cytometry analysis of SUM149 identifies that a large percentage of cells are CD24^hi^ CD44^hi^ (55), a proposed signature of the hybrid E/M phenotype (19).

JAG1 knockdown was effective in cell culture for up to 96 hours (Fig. 4A) but had no effect on viability or 2D proliferation in SUM149 cells (Fig. 4B-C). SUM149 cells, as an example of inflammatory breast cancer, are capable of spontaneously forming tumor 3D tumor spheroids or emboli when seeded in ultra-low attachment plates *in vitro* (57, 58). Thus, we utilized a tumor emboli in culture model allowing for the physiologically relevant 3D growth that is similar to the clinicopathological feature of collective tumor cell aggregates seen in patients with inflammatory breast cancer (57). Once the tumor emboli were formed at 48h, JAG1 siRNA treated emboli were significantly smaller and less dispersed that vehicle (p<0.001) or scrambled siRNA (p<0.0001) treated emboli (Fig. 4D-E). JAG1 siRNA treated emboli were harvested and lysed following 96 hours of growth, and found by immunoblot analysis to contain depleted levels of JAG1 and cleaved Notch proteins, as well as increased levels of DLL-4 protein, compared to vehicle treated cells (Fig. 4F). Overall, these findings indicate that the Notch-Jagged is a crucial axis that regulates the acquisition of proliferative traits.

**Figure 4.**
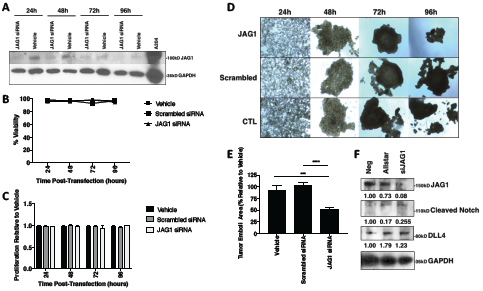
JAG1 knockdown reduces emboli size and decreases JAG1 pathway signaling. **A)** Western blot of SUM149 cells for JAG1 and GAPDH proteins following treatment with JAG1 siRNA or transfection reagent alone for 24, 48, 72, or 96h. **B)** Percent viability and **C)** 2D proliferation relative to vehicle of SUM149 cells at 24, 48, 72, and 96 hour timepoints following JAG1 siRNA, scrambled siRNA, or vehicle treatment. **D)** Representative 40x magnification images and **E)** area quantification of SUM149 tumor emboli treated with JAG1 siRNA, scrambled siRNA, or vehicle alone at days 1, 2, 3, and 4 post-transfection. **F)** western immunoblot analysis for JAG1, cleaved Notch, DLL4, and IL-6 proteins at t = 4 days (96h) from SUM149 tumor emboli lysates, normalized to GAPDH. ***p<0.001, ****p<0.0001 by one-way ANOVA and FLSD post-hoc test, n=6.

## Discussion

Targeting the ubiquitous intratumoral phenotypic and functional heterogeneity remains an unsolved clinical challenge. Such heterogeneity is exacerbated by non-cell autonomous effects of phenotypic plasticity. Phenotypic plasticity mediated by reversible activation of EMT or CSC pathways can fuel the acquisition of metastatically competent, drug refractory and adaptive phenotypes (22, 59). A canonical example of this heterogeneity is the presence of different subsets of CSCs with varying EMT phenotypes in spatially distinct regions of the tumor (9).

Here, using a mathematical modeling framework, we demonstrate that the spatial gradients of EMT-inducing signals such as TGF-β, coupled with cell-cell communication via Notch signaling, can generate distinct subsets of CSC with varying EMT phenotypes in spatially segregated regions of the tumor. Our model predicts that mesenchymal-like CSCs localize at the invasive edge of the tumor and hybrid E/M CSCs localize in the tumor interior. This prediction agrees well, at least qualitatively, with recent observations in human breast carcinoma tissue, where CD44+/CD24-cells (mesenchymal CSCs) were present at the tumor invasive edge, while ALDH1+ cells (hybrid E/M CSCs (20, 31)) were present in the interior of the tumor (9). However, we acknowledge that our model has multiple limitations. First, we assumed a spatial profile of TGF-β based on its secretion from stromal cells; however, there may be local variations in TGF-β signal intensity due to a variety of mechanical and/or chemical stimuli, including its secretion by cancer cells. Second, the model does not account for migration of hybrid E/M and mesenchymal cancer cells, which may modify the spatial patterning, particularly by overestimating the effects of mesenchymal cancer cells in inducing EMT in neighboring cells. Third, it considers EMT as a discrete three-step process, while recent studies point to a spectrum of phenotypic plasticity (60, 61). Despite these limitations, our framework offers promising novel insights into the spatial juxtaposition of different phenotypes within a given tumor tissue; such patterns are only beginning to be uncovered by advances in imaging techniques. Further progress in multiplexed imaging over large tumor sections can not only unravel the new layers of underlying biological complexity, but also present an opportunity to develop more representative and predictive computational models.

Our results highlight how inflammatory cytokines such as IL-6 can amplify Notch-Jagged signaling, thus facilitating cells to maintain a hybrid E/M stem-like phenotype, as well as form clusters of hybrid E/M cells that may dislodge from primary tumors as clusters of CTCs – the primary ‘villains’ of metastasis (16, 62). The presence of both the transmembrane receptor Notch and the transmembrane ligand Jagged can facilitate bidirectional signaling not only among cancer cells, but also in tumor-stroma crosstalk. For instance, in basal-like breast cancer (BLBC), where Notch, Jagged and IL-6 receptor (IL-R) are overexpressed relative to other breast cancer subtypes (10, 26), IL-6 upregulates JAG1, and boosts communication among cells through Notch3 and JAG1 (26). Notch3, a member of the Notch family, can facilitate the autocrine production of IL-6, hence ensuring sustained high levels of inflammatory cytokines (26). The IL-6/Notch3/JAG1 axis can sustain mammosphere growth; an autocrine activation of this axis seems to be required for higher invasive potential (26). These observations are reinforced by our results that JAG1 knockdown significantly impairs tumor formation potential in triple negative breast cancer (TNBC) SUM149 cells. Although not synonymous *per se*, TNBC and BLBC share numerous similarities, with over 70% overlap of TNBCs being classified as BLBCs, and *vice versa* (63). Furthermore, the association between Notch-Jagged signaling and hybrid E/M state is strengthened by observations such as a) loss of glycosyltransferase Fringe – a suppressor of Notch-Jagged signaling (64) – in BLBC (65), b) enrichment of hybrid E/M cells in primary tumor biopsies of TNBC patients (66), and c) association of enhanced JAG1 levels with poor outcome in BLBC (50). Beyond breast cancer, similar results for JAG1 knockdown are also seen in ovarian cancer, where silencing JAG1 in stromal cells as well as in tumor cells impaired tumor growth, and silencing it both in the tumor and stroma had synergistic effects (67). This synergistic effect can be explained by the observation that deleting JAG1 in stromal cells can limit the possibility of activation of Notch-Jagged signaling among tumor cells by limiting the strength of lateral induction, as noted experimentally (18).

Our results proposing that IL-6 driven Notch-Jagged signaling can mediate stemness are consistent with multiple previous studies in IBC and other subtypes. First, IBC emboli tend to have activated Notch3 intracellular domain; depletion of Notch 3 can lead to a loss of stem cell markers and induce apoptosis (68). Second, IBC patients have significantly higher levels of IL-6 levels in serum and carcinoma tissues as compared to non-IBC ones (69). Third, CD24^hi^ CD44^hi^ cells that represent a hybrid E/M phenotype (19) and form aggressive tumors *in vivo* (70) are enriched in IBC upon drug treatment (15) and exhibit upregulated Notch-Jagged signaling (5). Fourth, Notch-JAG1 signaling mediates resistance to tamoxifen (43), and is enriched upon inhibition of HER2 (41), suggesting how adaptive resistance mediated by JAG1 can contribute to tumor relapse. Put together, these studies strongly emphasize the role of inflammatory cytokines and Notch-Jagged signaling in cancer cell survival and aggressive potential.

## Methods

### Modeling the interplay between Notch signaling, EMT and inflammatory cytokines

The levels of every miR, TF and protein in the circuit is described by an ordinary differential equation that takes the form:

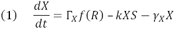

Where Γ*_X_* is the production rate, and *γ_X_* is the degradation rate constant, assuming first order degradation kinetics (see SI, sections 1, for the complete set of equations). Additionally, the presence of an activator/inhibitor (*R*) increases/decreases the basal production rate constant by a factor *f*(*R*). *f*(*R*) is a Hill function for transcriptional regulation, while it assumes a more complex form to describe protein degradation due to binding of non-coding RNAs to their target mRNAs (see SI sections 2). Additionally, Notch receptors and ligands irreversibly bind via a term – *kXS*, where *k* is a binding rate constant and *S* is the receptor/ligand binding to *X*. This term is present only in equations where *X* is a Notch receptor/ligand. Binding affinity of Notch receptor is different for Delta and Jagged ligands. Additionally, ligands can inhibit the signaling by binding to Notch receptors of the same cell which leads to the degradation of the complex (cis-inhibition). In this scenario, both *X* and *S* belong to the same cell.

In the model, inflammatory cytokines such as TNF-α, IL-6 and IL-1β are described by a variable *C_EXT_*, which represent the number of inflammatory molecules that interact with the cell, inhibiting the production of Delta and activating Jagged according to:

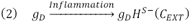

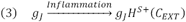

Where *H^S-^*/*H^S+^* are negative and positive shifted Hill functions, respectively (37).

This framework is extended to model the dynamics of a two-dimensional hexagonal lattice of cells. In the multi-cell model, Delta and Jagged ligands of a cell can bind to the Notch receptors of a neighbor cell. Therefore, the number of receptors and ligands available to bind to one cell’s receptors and ligands are:

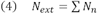

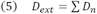

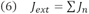

where *N*_n_, *D*_n_, *J*_n_ are the levels of Notch, Delta and Jagged in the neighbor cells n, and the index n represents the set of nearest neighbors in the lattice.

The cell tissue can be exposed to an EMT-inducing signal *I*(*x*, *y*, *t*) such as TGF-β that is secreted at one end of the lattice and removed at the opposite end (a source-sink dynamics). This profile mimics the situation that this diffusible molecule can be secreted by many stromal cells that are present next to the tumor invasive edge. Therefore, this signal obeys a diffusion equation:

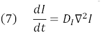

with boundary conditions such that *I* = *I_max_* at the tissue end that secrets the signal (*x* = 0) and *I* = 0 at *x* = *L*, where *L* is the length of the tissue.

### Cell culture and reagents

SUM149 (Asterland, Inc.) inflammatory breast cancer cells were cultured as per manufacturer’s instructions and previous studies (71). Cells were cultured with 1% penicillin/streptomycin (Invitrogen) supplemented in their respective media. Cells were cultured in growth medium at 37°C under an atmosphere of 5% CO2.

### Cell transfection

Cells were plated at 20,000 cells/well in a 24-well plate and transfected 24h later with 37.5ng siJAG1 (QIAGEN#SI02780134), Allstar scrambled siRNA (QIAGEN #SI03650318), or no siRNA with HiPerfect (QIAGEN #301704) transfection reagent added alone. siRNA was incubated with 3ul of transfection reagent at room temperature for 10 minutes and added drop-wise to each respective well on the plate. Cells were incubated at 37C.

### Trypan Blue exclusion viability assay analysis

Cells were plated in a 12-well plate at 3×10^4 cells per well for 24h, then treated with PAH mixture, chlorothalonil, or left untreated. Cells were trypsinized after 24h and resuspended in media, then centrifuged at 1600rpm for four minutes. The supernatant was aspirated and 70ul cold DPBS added to the pellet. An aliquot of cell suspension was mixed with an equal volume of 0.4% trypan blue solution, and cell numbers were recorded in 10ul of the resultant mixture using a hemocytometer. Viability is reported as the ratio of live cells to total cells.

### MTT proliferation assay analysis

Cell toxicity and proliferation were assessed by MTT assay (3-[4,5-dimethylthiazol-2-yl]-2,5 diphenyl tetrazolium bromide), based on the conversion of MTT into formazan crystals by living cells, which determines metabolic activity and is an acceptable measure of viable, proliferating cells. Cells were seeded at 4,000 cells/well into 96-well flat bottom plates and transfected. Cells were grown for 24, 48, 72, or 96 hours. Proliferation was assessed using 3(4,5-dimethylthlthiazol-2-yl)-2,5-diphenyltetrazolium, after which cells were incubated at 37°C for 2h, DMSO was added to each well, and absorbance was read at 570nm on a Fluostar Optima plate reader.

### Tumor organoid formation analysis

SUM149 cells were plated at 10,000 cells per well in an ultra-low attachment 24-well plate with filtered media supplemented with 2.25% polyethylene glycol as described previously (57). The resultant tumor cell emboli in culture were left to form for 48 hours, verified formed and uniform by microscopy, then assessed for growth and size. Tumor emboli in culture were then allowed to grow for 72-96 hours, after which they were imaged at 4x magnification and their gross particle area was analyzed using NIH Image J.

### Western immunoblot analysis

Tumor emboli in culture were harvested after 96h growth and lysed, and immunoblot analysis was performed as described previously (72). 2D monolayer cells were also treated and harvested following transfection for further immunoblot studies. Membranes were incubated overnight at 4°C with primary antibodies JAG1, cleaved Notch, DLL4, IL-6 (Cell Signaling Technologies, 1:1000 dilution), or GAPDH (1:2000 dilution). Membranes were washed and incubated with anti-mouse or anti-rabbit HRP-conjugated antibodies (Cell Signaling Technologies) for 1 hour at room temperature. Chemiluminescent substrate was applied for 5 min, and then membranes were exposed to radiographic film. Densitometric analysis was performed using NIH ImageJ software with GAPDH as a loading control, with all values normalized to the untreated lanes.

## Author contribution

Conceived research: Mohit Kumar Jolly, Marcelo Boareto, Eshel Ben-Jacob; Mathematical model construction: Federico Bocci, Marcelo Boareto, Mohit Kumar Jolly; Numerical calculations: Federico Bocci; JAG1 knockdown: Larisa Gearhart-Serna, Mariana Ribeiro; Writing: Federico Bocci and Mohit Kumar Jolly. All authors discussed the results and edited the manuscript; Guided research: Gayathri Devi, Herbert Levine, José Nelson Onuchic, Mohit Kumar Jolly.

## Acknowledgements

This work was sponsored by the National Science Foundation (NSF) grants PHY-1427654 (Center for Theoretical Biological Physics), PHY-1605817, CHE-1614101, MCB-1241332 and by the Cancer Prevention and Research Institute of Texas (CPRIT) grant R1110 and in part by Department of Defense grant W81XWH-17-1-0297 (GRD) and Duke School of Medicine Research Funds (GRD). Larisa Gearhart-Serna was supported by NIEHS T32-ES021432. Federico Bocci was partially supported by the Hasselman Fellowship for academic excellence in Chemistry. Mohit Kumar Jolly was also supported by a training fellowship from the Gulf Coast Consortia, on the Computational Cancer Biology Training Program (CPRIT Grant No. RP170593). Marcelo Boareto was supported by FAPESP Grant 2013/14438-8

## Additional information

Competing financial interests: the authors declare no competing financial interests.

